# Delayed post-juvenile moult in malaria-infected European blackcaps

**DOI:** 10.1101/2024.07.31.605960

**Authors:** Carolina Remacha, Iván de la Hera, Álvaro Ramírez, Javier Pérez-Tris

**Affiliations:** Department of Biodiversity, Ecology and Evolution, Complutense University of Madrid, Madrid, Spain

**Author notes:** **Correspondence:** Carolina Remacha.

**Keywords:** post-juvenile moult, haemosporidian parasites, *Sylvia atricapilla*, parasitemia

## Abstract

Parasites have their strongest impact on fitness when host defences limit the quantity of resources available for other critical life-history stages, such as development, breeding or migration. One greatly neglected stage that could be altered by parasites is the post-juvenile moult (PJM) of birds, through which inexperienced yearlings replace their weak and fast-generated juvenile feathers by adult-like feathers. The earlier juvenile birds complete PJM, the earlier they will be prepared to withstand forthcoming adverse conditions, especially if they migrate short after moulting. We used data from 435 juvenile European blackcaps (*Sylvia atricapilla*) sampled during three years in 26 localities spanning the wide range of environmental conditions across Iberian Spain to test whether haemosporidian infections (presence and abundance in blood of parasites of the genera *Haemoproteus, Plasmodium* and *Leucocytozoon*) were related to delayed PJM. Controlling for body condition, sex, year and date of capture, infected blackcaps (single-infected or coinfected) had lower moult scores indicative of delayed moult, especially when birds had *Plasmodium* infections or higher intensity of *Haemoproteus* parasites. Our results broaden the range of fitness costs that haemosporidian parasites may have on birds, as delayed plumage maturation prolongs the period during which juvenile birds lack a fully functional plumage.

## INTRODUCTION

Parasitised animals need to allocate body resources to self-defence that otherwise would be available for other demanding activities, such as development, reproduction, or migration [1]. When infections are not lethal, parasites can still considerably lower host fitness by reducing reproductive success or lifetime expectancy [2,3]. Understanding parasite impacts requires uncovering the physiological mechanisms through which parasite exploitation impairs host performance during life stages that are critical to maximize fitness. These stages include not only episodes of breeding investment or periods when survival may be compromised (e.g. times of food shortage or energy-demanding activities such as migrations), but also periods of development when phenotypic traits are produced that may determine breeding or survival later in life. A textbook example are bird parasites that impair the development of bright plumage ornaments [4], an impact that necessarily takes place during feather moult but often has its consequences on the bird’s fitness several months later, during the mating season when feather ornaments function as honest signals of individual quality [5].

Moult is an energetically demanding stage in the life cycle of birds, as feathers may account for almost a quarter of the total protein content of a bird’s body [6]. Apart from other auxiliary roles [7], moult plays two main non-exclusive functions in birds. First, by replacing old feathers by new ones, it restores plumage functionality. Second, moult also evolved to adjust plumage characteristics (e.g. coloration, feather structure) to new requirements birds meet as they age or change environmental conditions [8].

However, the costs of feather production lead to trade-offs between moult and other vital functions such as stress response [9], oxidative balance [10], or immunity [11,12]. An activation of the immune system can delay the onset of moult [13], and individuals with parasites may develop worse plumage [14,15]. These adjustments are related to metabolite competition [7] and interference with biochemical processes, such as corticosterone negatively influencing melanogenesis and varying seasonally with moult or condition [16,17].

During feather shedding, thermoregulatory and flight functions of the plumage are impaired [8,18] so individuals try to avoid overlapping moult with other demanding activities, such as breeding or migration [19]. Nestlings of most passerine species develop quickly producing low quality juvenile feathers that allow them to abandon the nest as soon as possible to shorten the period of high risk of predation [20]. Soon after fledging, juvenile birds need to replace part or the whole set of juvenile feathers by new ones [21] to improve flight efficiency during migratory travels and/or thermal properties to withstand adverse thermal conditions [22]. Known as the post-juvenile moult (hereafter PJM), this maturation process has been often overlooked despite its strong consequences for bird fitness at this vulnerable age [23]. The timing of PJM is crucial because the sooner it is completed, the sooner juvenile birds will be better prepared to face adversity (Morales et al 2007). Interestingly, parasites may have an impact on bird fitness if parasitised individuals have an impaired performance during this critical life-history event. Haemosporidian parasites are widespread vector-borne blood parasites commonly found in many bird species [25] for which there is ample evidence of their adverse effects on host fitness and survival [26]. The life-cycle of these parasites in bird hosts includes different stages of development, first reaching some tissues before their spreading into the bloodstream to invade mainly erythrocytes [27]. Some infections remain chronic with seasonal relapses matching periods of highest probability of transmission that combine high vector availability and low host coping ability [27]. The virulence of infections depends largely on the combination of parasite type and bird species [28,29] and impacts can become more evident when resources are compromised due to trade-offs with other demanding vital processes [30].

Plumage functions such as honest signalling by feather coloration seem to be compromised by these parasites [31,32]. However, the few studies attempting to address the influence of haemosporidian parasites on bird moult progress have not achieved clear conclusions [24,33] and often were based on mismatched parasite loads estimated outside the moulting period. Furthermore, infected birds seem to replace feathers more slowly [34,35], but these results have been based on experiments inducing feather replacement, which can produce confusing results because the cost of growing feathers outside the moulting season can be low or differ seasonally [36,37]. Therefore, it is especially important to study the influence of infection status on moult when the process is naturally occurring.

Our aim was to test whether infection by haemosporidian parasites was related to PJM progress in first-year Eurasian blackcaps (*Sylvia atricapilla*) from several populations distributed throughout the Spanish part of the Iberian Peninsula. This species faces high haemosporidian prevalence soon after fledging [38] and juveniles perform a partial PJM before their first migration [39]. Co-infections by different parasites and parasitemia (abundance of parasites in blood) were considered in the analyses as they can sometimes modulate the impact of infections [40–42]. We also tested if the influence of infection on moult progress could be different in males and females because sex hormones are important modulators of the immune response [43]. Finally, we controlled for the influence of body condition and date of capture in the models because timing of PJM is sensitive to seasonal changes in photoperiod [44,45] and individual quality [46,47]. We expect infected blackcaps to have delayed PJM, more so in birds with higher parasite load or in coinfected individuals.

## MATERIAL AND METHODS

### Bird species and sampling

The Eurasian blackcap is a small passerine bird widely distributed throughout the Western Palaearctic and Macaronesia with most mainland populations migratory [48]. Short after fledgling (44-47 days old, [49]), juveniles undergo a partial PJM typically involving most wing coverts and all body contour feathers, including ornamental traits such as the sexually dimorphic crown (black in males and brown in females). However, most flight-feathers remain un-moulted until the first complete moult the following year [49,50]. The PJM lasts around 44 days in blackcaps [49].

Thirty-three populations of blackcaps distributed across Iberian Spain (Figure 1) were sampled in 2008, 2009 and 2011 during the post-fledging period (July and August) before migration started. For this study we only used data of 26 localities where at least 10 juveniles were captured. These sampling sites represent a wide range of environmental and landscape features (altitude, slope, percentage of broadleaved forest, wooded croplands, arable lands or urban areas), spanning the range of habitat suitability for Iberian blackcaps (see full sampling details in [51]). Birds were captured with mist nets and conspecific song playbacks during the morning and the afternoon. Individuals were marked with official aluminium rings and measured tarsus length (0.01-mm precision) and body mass (to the nearest 0.01 g) among other traits recorded during their handling. We scored the stage of their PJM using a scale that ranges from 0 (juvenile primary feathers still growing up and moult not yet started) to 6 (post-juvenile moult completed), based on the pattern of moult of specific feather tracts in the throat, flanks and belly [52]. PJM scores were taken by two observers (IH, JP-T) and there were no significant differences between them (F1,422.2 = 2.44, *P* = 0.12, linear mixed model). One blood sample was obtained through brachial or jugular vein puncture according to safety protocol ([53], < 1% body mass). First, a drop of blood was spread on a slide for microscopy analyses, and remnant blood was stored in absolute ethanol for molecular analyses.

**Figure 1.**
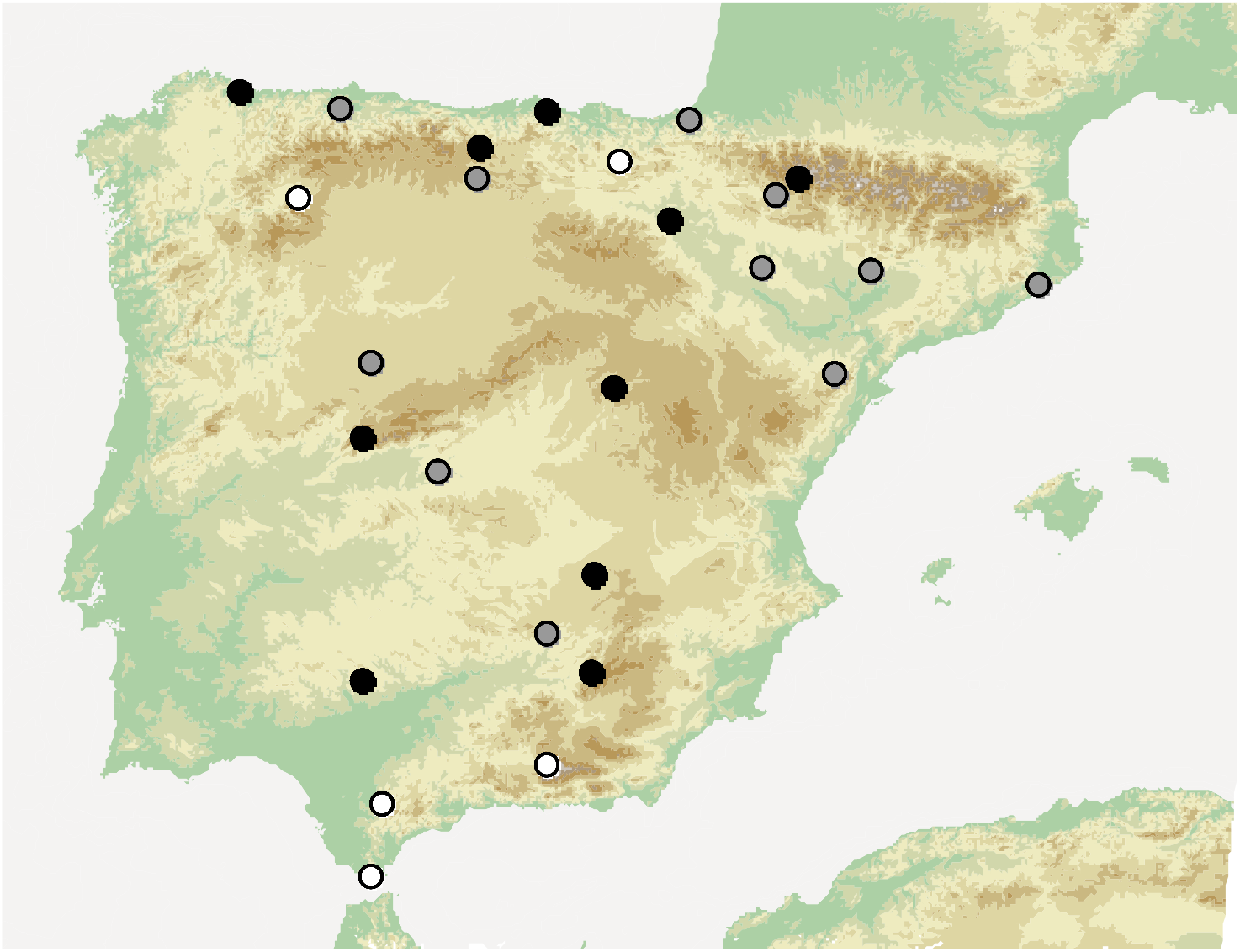
Map of the Iberian Peninsula with sampling locations where at least 10 juveniles were captured in 2008 (white), 2009 (grey) or 2011(black circles).

### Parasite assessment

DNA was extracted with standard ammonium acetate protocols [54], quantified and diluted to a working concentration of 25 ng/μl. Previous to molecular parasite screening we sexed birds using a PCR-based protocol ([55]; young blackcaps cannot be sexed in the field before they have moulted crown feathers), which also served as a PCR-quality control of samples.

Parasites were detected using a nested method that amplifies a 478-bp fragment of the *cytochrome b* mitochondrial gene in two steps [56]. The first PCR includes primers (HAEMNFI and HAEMNR3) and amplifies a longer fragment including *Haemoproteus, Plasmodium and Leucocytozoon* DNA. Conditions were as following: 1 μl of each 10 μM primer, 2.5 μl of a mix of 1.25 mM of each dNTP, 1.1 μl of 25 mM of MgCl2, 2.5 μl of buffer 10×, 14.8 μl of ddH2O and 2 μl of DNA template. The thermal profile included a denaturation step at 94 °C for 3 min; 20 cycles of 94 °C for 30 s, 50°C for 30 s and 72 °C for 45 s, and an extension of 10 min at 72 °C. Two downstream specific PCRs included 1μl of the previous PCR product as the template with the same conditions but increasing the number of cycles to 35. PCR with HAEMF and HAEMR2 primers targeted *Haemoproteus* or/and *Plasmodium* DNA, and PCR with HAEMFL and HAEMR2L primers targeted *Leucocytozoon* DNA. Positive samples were sequenced on an ABI Prism 3730 capillary robot (Applied Biosystems, Paisley, UK) and aligned using software BioEdit 7.0.5.3 [57]. Following the standards of the MalAvi database, sequences differing in one single nucleotides were deemed different lineages [25]. In this way, birds harbouring more than one lineage were considered coinfected.

Parasite load was assessed by counting the number of infected erythrocytes in 10000 in Giemsa-stained blood smears with a LEICA DM2500 microscopy (Leica Microsystems, Wetzlar, Germany).

### Statistical analyses

The relationships between PJM and infection were assessed with Bayesian ordinal mixed regression models using the *brms* package [58]. Models were fitted using a sequential distribution and *logit* link, because PJM categories describe a sequential process according to which the birds that reach a higher moult value have necessarily gone through the previous moult stages [59]. We included infection, sex, and their interaction when it was significant as fixed effects. Year, body condition and Julian date of capture were also included as covariates, and locality as a random factor. Marcov-Chain Monte Carlo (MCMC) simulations were run using four chains for each model with 110000 total iterations (the first 10000 iterations were discarded as warmup), which were sampled with a thinning interval of 100 iterations. We assessed model convergence with posterior predictive checks (*pp_check* function from the bayesplot package), together with R-hat, Bulk_ ESS (the effective sample size for rank normalized values using split chains) and Tail_ESS values (the minimum of the effective sample sizes for 5% and 95% quantiles), and plots of the posterior density and trace values for each parameter [60,61]. We tested the influence of infection using several complementary approaches. We first considered the multiple status of infection of individuals (uninfected, single-infected or coinfected) irrespective to the identity of the parasites involved). Secondly, we considered the status of infection by each genus (*Plasmodium, Haemoproteus* and *Leucocytozoon*) separately. We also tested the influence of the parasite load, total and for each genus, including only the subset of infected birds diagnosed either by PCR or microscopy (not infected birds cannot be quantified a parasite intensity, so that zero-intensity infections are those infected birds for which no parasite was found in counts of 10000 erythrocytes). *Plasmodium* and *Leucocytozoon* gametocyte presence was too low to measure their parasitemia on a continuous scale, so we classified their parasite load in two categories: presence or absence of blood gametocytes. In addition, models analysing *Plasmodium* and *Leucocytozoon* parasite load with all covariates showed signs of overfitting, which was solved by excluding the year factor.

Frequency of some PJM categories was too low (three birds with PJM score 0 and two with score 5), so we repeated the analyses removing these data and reported the corresponding results in the main text when there was a discrepancy with the models that included all data (see all results in electronic supplementary material B).

Body condition was calculated as the residuals of the body mass against tarsus length (estimate = 0.52, F1,432 =47.26, *P* < 0.001, [62] and time of capture measured as a dichotomous factor (morning or afternoon given the bimodal distribution of sampling hours; estimate = -0.97, F1,432 = 83.10, *P* < 0.001). All analyses were performed with R software (version 4.3.0) and additional packages (see complete list and references in available R code). Continuous variables were standardized by subtracting the mean from each value and dividing by the standard deviation before running each model.

Residual distributions and other model assumptions were checked in all analyses. Sample size differed among analyses due to missing values, for example in 2011 we have no data for parasite intensity for 6 localities.

## RESULTS

Observed PJM scores ranged between 0 and 5 (mean ± sd = 1.79 ± 1.00; Figure 2). Mean haemosporidian (n = 435 birds) prevalence was 70.17 ± 28.49 % (16.20 ± 19.38% for *Plasmodium*, 61.06 ± 34.34 % for *Haemoproteus* and 10.33 ± 17.82 % for *Leucocytozoon*). In almost all our models PJM was more advanced in females compared to males, and it was positively correlated with individual body condition (estimates for these variables in each model are shown in electronic supplementary material A S1-S8, not significant in the model analysing *Leucocytozoon* parasite load).

**Figure 2.**
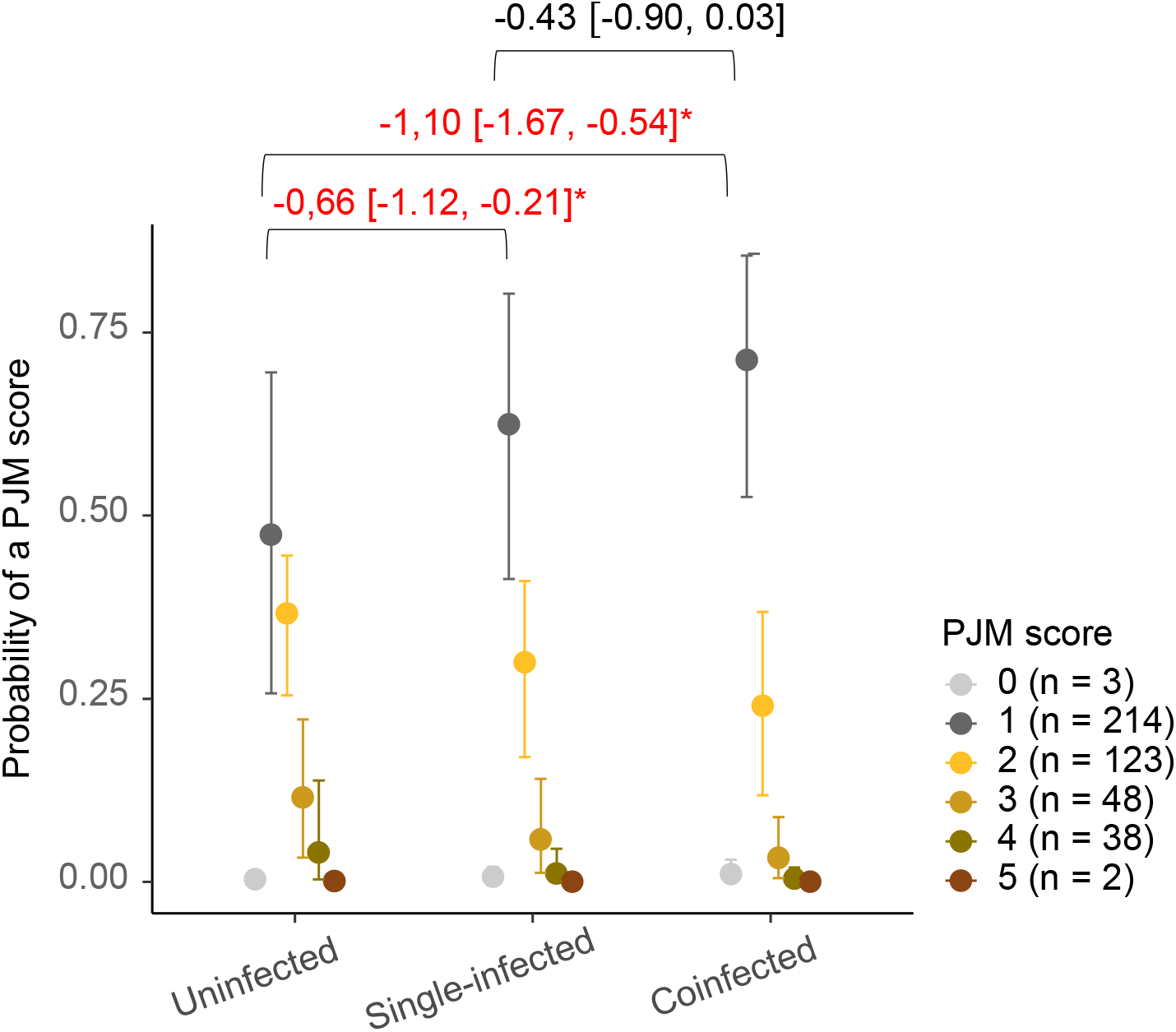
Mean posterior probability and 95% Bayesian credible intervals (BCI) of each PJM score (represented with different colours) in juvenile blackcaps that were either uninfected, single-infected or coinfected by any haemosporidian lineage. Bayesian models controlled for sex, body condition, year, and date of capture. The top part of the figure shows average posterior predicted estimates and 95% BCI for pairwise comparisons of infection levels. Comparisons with 95% BCI not overlapping zero were deemed statistically significant and are highlighted using red colour and asterisks. Sample sizes of each PJM score are indicated in the legend (seven birds that could not be resolved as single-infected or coinfected were not included in this analysis).

Uninfected birds were more advanced in their PJM compared to birds with either single infections or coinfections (Figure 2) after controlling for sex, body condition, year, and date of capture (Electronic supplementary material A Figure S1). Differences between single-infected and coinfected birds were close to statistical significance (Figure 2).

Blackcaps with *Plasmodium* infections had a delayed moult compared to birds without infections of this genus (estimate = -1.08, 95 % BCI = [-1.64, -0.53]) after controlling for sex, body condition, year, and date of capture (Electronic supplementary material A Figure S2). *Haemoproteus*-infected birds were also delayed in their PJM (estimate = -0.44, 95 % BCI = [-0.88, -0.02]) after controlling for sex, body condition, year and date of capture (Electronic supplementary material A Figure S3). *Leucocytozoon* infection was not significantly correlated with PJM (estimate = 0.42, 95% BCI = [-0.27, 1.12]; see all covariate effects in Electronic supplementary material Figure S4).

Total parasite load of infected birds was weakly correlated with PJM (estimate = -0.32, 95 % BCI = [-0.68, 0.03]), a weak effect which vanished when infrequent PJM scores were excluded from the analysis (estimate = -0.26, 95 % BIC = [-0.64, 0.09]; see all covariate effects in Electronic supplementary material Figure S5). PJM was not significantly correlated with presence of *Plasmodium* (estimate = -0.80, 95 % BCI = [-2.25, 0.54]) or *Leucocytozoon* gametocytes (estimate = -0.60, 95 % BCI = [-2.44, 1.09]; see all covariate effects in Electronic supplementary material Figure S6 and S8 respectively), but *Haemoproteus* parasite load was negatively correlated with PJM scores (estimate = -0.52, 95 % BCI = [-0.95, -0.11], Figure 3), although the effect was weaker when infrequent PJM scores were excluded (estimate = -0.43, 95 % BCI = [-0.88, 0.02]; see covariate effects in Electronic supplementary material A Figure S7). Most clearly, a higher intensity of *Haemoproteus* infection was associated with a lower probability of having transitioned from PJM score 2 to 3 (Figure 3).

**Figure 3.**
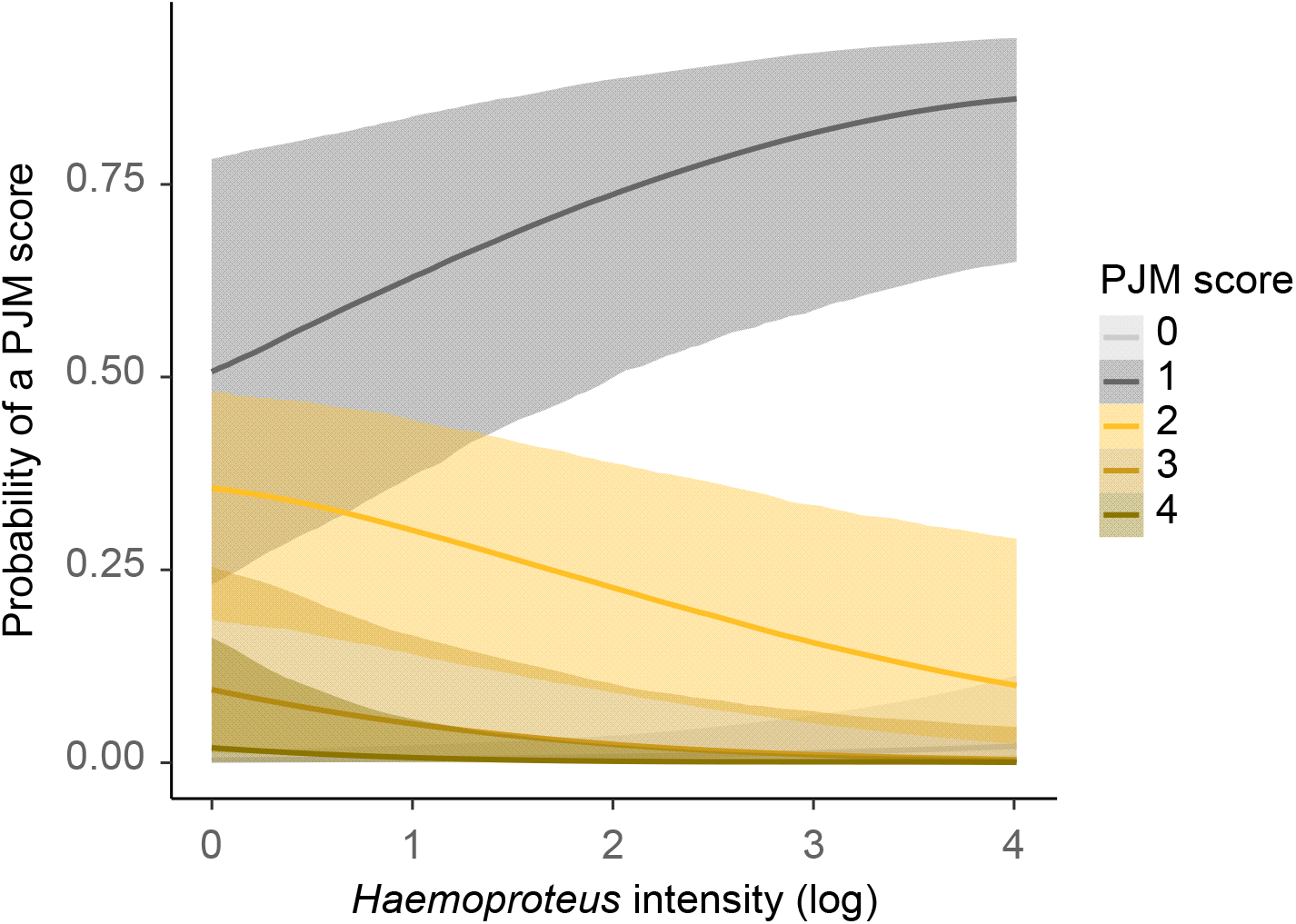
Relationships between *Haemoproteus* intensity (log scale) and the posterior probability of each PJM score (represented with different colours) in juvenile blackcaps in Bayesian models controlling for sex, body condition, year, and date of capture. Shaded areas represent 95% Bayesian credible intervals (BCI) of each estimated relationship (lines).

## DISCUSION

Infected blackcaps had delayed post-juvenile moult. When the correlation between infection status and body moult score was analysed separately for each parasite genus, *Haemoproteus* and (especially) *Plasmodium* showed the strongest effects. *Leucocytozoon* parasites had a low prevalence and were not clearly associated with moult stage. We found weak evidence that coinfected birds had delayed moult compared to single-infected individuals. In addition, *Haemoproteus* parasite load was related with delayed post-juvenile moult, although this effect was weaker when infrequent moult scores were excluded from its analysis. Finally, females and birds in better body condition had a more advanced moult. Given the geographic extent of our sampling, these results can be safely taken as general to the range of environmental conditions encountered by Iberian Spanish blackcap populations.

Our results suggest that parasite effects on PJM vary among parasite genera. The presence of *Plasmodium* was associated with delayed PJM despite the low intensity of infections of this parasite genus (in only 15 individuals, parasites were observed by microscopy). For *Haemoproteus* parasites, which showed a broader range of infection intensities than *Plasmodium*, we found effects of both individual status of infection and infection intensity, as infected birds or those with more gametocytes had delayed PJM. Among the few studies addressing the relationships between haemosporidian infections and feather growth, Marzal et al. [41] found on house martins *Delichon urbicum* that coinfections decrease feather growth, which could be related to higher parasite loads if coinfections involve higher parasitaemia [63]. Our results suggest that not only intensity of infection, but also parasite identity, may be important determinants of the impact of parasites on the PJM process, reinforcing the idea that *Plasmodium* parasites are especially virulent to blackcaps. *Plasmodium relictum*, the most frequent lineage of this genus in our population [64], is a widespread generalist parasite which has been shown to strongly impact on host health and survival [65,66], although its virulence might vary among species [65,67,68] or even among individuals of the same species [69]. Experimental infections with the *Plasmodium relictum* lineage SGS1 may reduce feather growth rate as found in house sparrows *Passer domesticus* [35]. Our interpretation that *Plasmodium* infections may be especially virulent to moulting blackcaps too is coherent with other studies showing negative relationships between *Plasmodium* infections and diverse indicators of host performance in this species, regardless of the parasitemia [70,71].

Haemosporidian parasite intensity and IgY levels have been shown to be positively correlated in juvenile blackcaps [72], which supports the interpretation that an activation of immune defence would delay moult if both processes compete for limited resources. Induced immune challenges have been found to impair feather growth in several bird species suggesting a trade-off between both processes [12,13], which may be mediated by limiting metabolites such as cysteine. This amino acid is implied both in the production of glutathione (GSH), an antioxidant that controls host damage caused by the activation of immune defences against parasites, and in the production of keratin, the main component of feathers [73]. Cecere et al. (2016) also found in juvenile European blackbirds *Turdus merula*, that moulting individuals showed higher oxidative damage, while an increased intake of dietary antioxidants could fasten their PJM. Therefore, birds with parasites may need to allocate cysteine to maintaining oxidative balance, at the expense of slow feather production, delaying moult completion as a consequence. This relationship may be exacerbated by the fact that haemosporidian parasites obtain essential amino acids from their host, which include isoleucine, leucine or valine, resources that are also necessary for the production of feathers [74–76].

A delayed moult can be especially deleterious for migratory birds, such as blackcaps from most populations. Moult strategies have evolved to avoid overlapping moult with other energy demanding processes such as breeding or migration, thus moulting late may have negative consequences on subsequent migration and juvenile dispersal [8]. The end of PJM has been related with the starting of autumn migration in blackcaps [22]. Thus, if haemosporidian infections delay moult, the migratory schedule of infected individuals could also be altered. Late migration may be disadvantageous if birds arriving early at their wintering ground monopolise the best habitat patches and/or promote social dominance over late-arriving individuals [77,78]. Furthermore, timing of arrival to non-breeding grounds may have carry-over effects on fitness if birds occupying the worst wintering sites delay the onset of spring migration, which is correlated with lower breeding success in many species [79–81]. Additional constraints caused by parasites during migration are especially important for young birds, which due to their lower social status and inexperience tend to arrive later and attain lower breeding success than adults [79]. Nevertheless, we do not know the exact date in which each juvenile completed moult and since the rate of PJM can vary among individuals (the onset of moult was not correlated with the end of moult in blackcaps; [22]), further longitudinal studies would be necessary to confirm an interference of parasites with migration via delayed moult. Anyway, if infected birds were able to accelerate their moult to compensate for a bad start, it would likely be at the expense of a poorer quality of the plumage [82], with similar consequences to those described above.

Not knowing the hatching date of individuals is a limitation in our study, as PJM score has been used as a proxy of time since hatching in small passerines [52,83]. If this was the case in Iberian blackcaps, our results would be best interpreted as late-hatched, younger birds being more likely infected, which would be possible if late broods were either more exposed to vector activity or attained lower body condition and reduced immunocompetence [84]. Nevertheless, vector activity is high in Iberian forests long before blackcaps start brooding (typically in May [85]), and we controlled body condition and date of capture in our analyses, which minimised the influence of phenotypic quality and phenology. In addition, the relationship between age and moult progress has been found at high latitudes where the breeding-moulting season is much shorter, and therefore this relationship may be tighter. For example, willow warblers in Swedish Lapland initiate PJM at age 26 days, right after outer primaries attain full length, and take around 18 days to reach score 5 (moult nearly completed; [52]). However, various lines of evidence suggest that the relationship between age and moult score may be weak in young Spanish blackcaps. The duration of the post-juvenile moult increases in blackcap populations found at lower latitudes (Figure 9 in [86]). In addition, late-hatched blackcaps can have a faster moult than early-hatched ones [87]. Therefore, we are confident that our results are not driven by age differences, and rather reflect an impact of parasite infection on the moult process.

Interestingly, PJM was more advanced in females than in males, but this difference did not mediate the relationship between infection and moult. Influence of sex hormones on immune responses fluctuate seasonally, and the strongest effect has been found during adults’ breeding or during the development of sexual ornamental traits [88]. Nevertheless, blackcaps develop their sexually dimorphic crown feathers at the end of PJM. Lastly, males frequently have delayed PJM, a pattern which has been attributed to males usually having more extensive moults than females [50,83].

Positive associations between body condition and moult progress have been previously found [89] and several mechanisms might drive this relationship. On the one hand, higher availability of nutrients in birds with better body condition could contribute to a fast moulting [90]. Alternatively, the observed correlation could be also due to physiological changes necessary for feather shedding in which individuals store water and other nutrients in the growing feathers that can increase their weight [91].

In summary, our results contribute to improve our understanding of how haemosporidian infections can impair bird performance, providing further hints that their effect is dependent on parasite genus and infection intensity. By adding post-juvenile moult to the list of critical life-history stages these parasites may interfere with, our results broaden the knowledge of potential parasite impacts on bird fitness, either directly or as a consequence of negative carry-over effects of impaired development.

## Supporting information

Electronic Supplementary Material A

Electronic Supplementary Material B

## ETHICS

The Spanish Regional Governments authorized fieldwork, with the following permits: Diputación Foral de Álava (Exp 08/32), Generalitat de Catalunya (SF/530-535), Generalitat Valenciana (2009/23822), Gobierno de Aragón (LC/ehv 24/2011/3629; LC/mp 24/2009/4370), Gobierno de Cantabria (SEP 257/11), Gobierno de La Rioja (A/2011/40/LL/aic), Gobierno de Navarra (SCB/20090524), Gobierno del Principado de Asturias (2009/024357), Junta de Andalucía (SGYB/FOA/AFR/MVAC; SGYB-AFR-CMM), Junta de Castilla y León (IS/pa/EP/SG/374/2008; IS/rp/EP/SG/53/2009), Junta de Comunidades de Castilla-La Mancha (DGAPYB/SB/avp/11_187; OAEENN/SVS/cgs), Junta de Extremadura (LLM/jcm/CN0041/11/AAN), and Xunta de Galicia (71/2011).

## DATA ACCESSIBILITY

Data and scripts are accessible from the corresponding author upon reasonable request.

## DECLARATION OF AI USE

We have not used AI-assisted technologies in creating this article.

## AUTHORS’ CONTRIBUTIONS

CR: Conceptualization; Data curation; Formal analysis; Investigation; Methodology; Visualization; Writing – original draft; Writing – review & editing

IH: Investigation; Methodology; Writing – review & editing

AR: Investigation; Methodology; Writing – review & editing

JP-T: Conceptualization; Data curation; Supervision; Funding acquisition; Investigation; Methodology; Project administration; Resources; Writing – review & editing

All authors gave final approval for publication and agreed to be held accountable for the work performed therein.

## ACKNOWLEDGEMENTS

We are indebted to Antón Pérez-Rodríguez and Sofía Fernández-González, who greatly contributed to generate this database during their PhD. Both assisted during fieldwork sessions, and APR did all molecular and microscopy analyses. We thank Roberto Carbonell for help during fieldwork.

## FUNDING

This study was funded by MCIN/AEI/10.13039/501100011033 and “ERDF A way of making Europe” through grants CGL2007-62937/BOS, CGL2010-15734/BOS and PID2020-116121GB-I00) and the Department of Education, Universities and Research of the Basque Government (studentships BFI. 04-33 and 09-13, to IH).

## Notes

### Competing Interest Statement

The authors have declared no competing interest.

