## Supplementary material for "Delayed post-juvenile moult in malaria-infected European blackcaps": Electronic Supplementary Material A

Electronic supplementary material A S1. Posterior distributions with medians and 95% confidence intervals for each parameter of the bayesian model assesing the influence of multiple status of infection (uninfected, simple infected and multiple infected) on post-juvenile moult, controlling for year, sex, body condition and date of capture. We used * for significant comparisons with 95% BCI range values not overlapping zero.

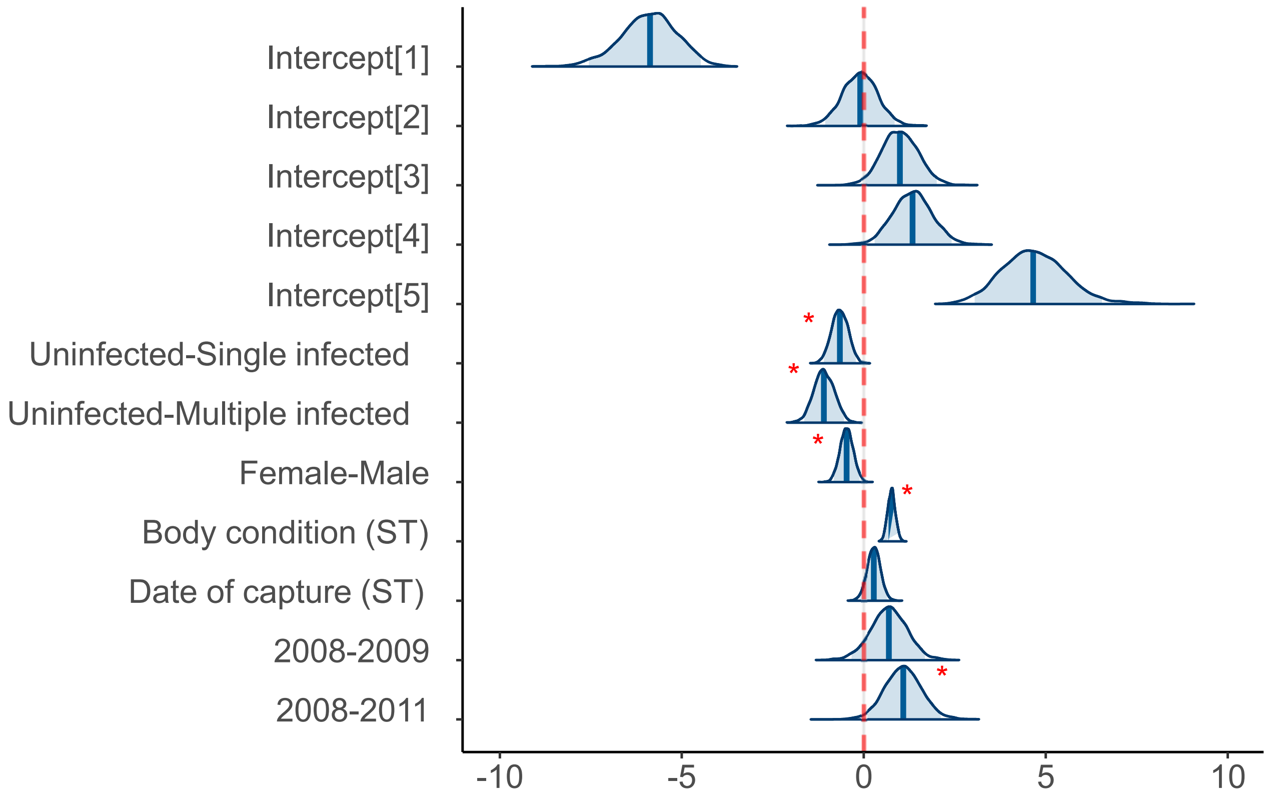

Electronic supplementary material A S2. Posterior distributions with medians and 95% confidence intervals for each parameter of the bayesian model assesing the influence of *Plasmodium* status of infection on post-juvenile moult, controlling for year, sex, body condition and date of capture. We used * for significant comparisons with 95% BCI range values not overlapping zero.

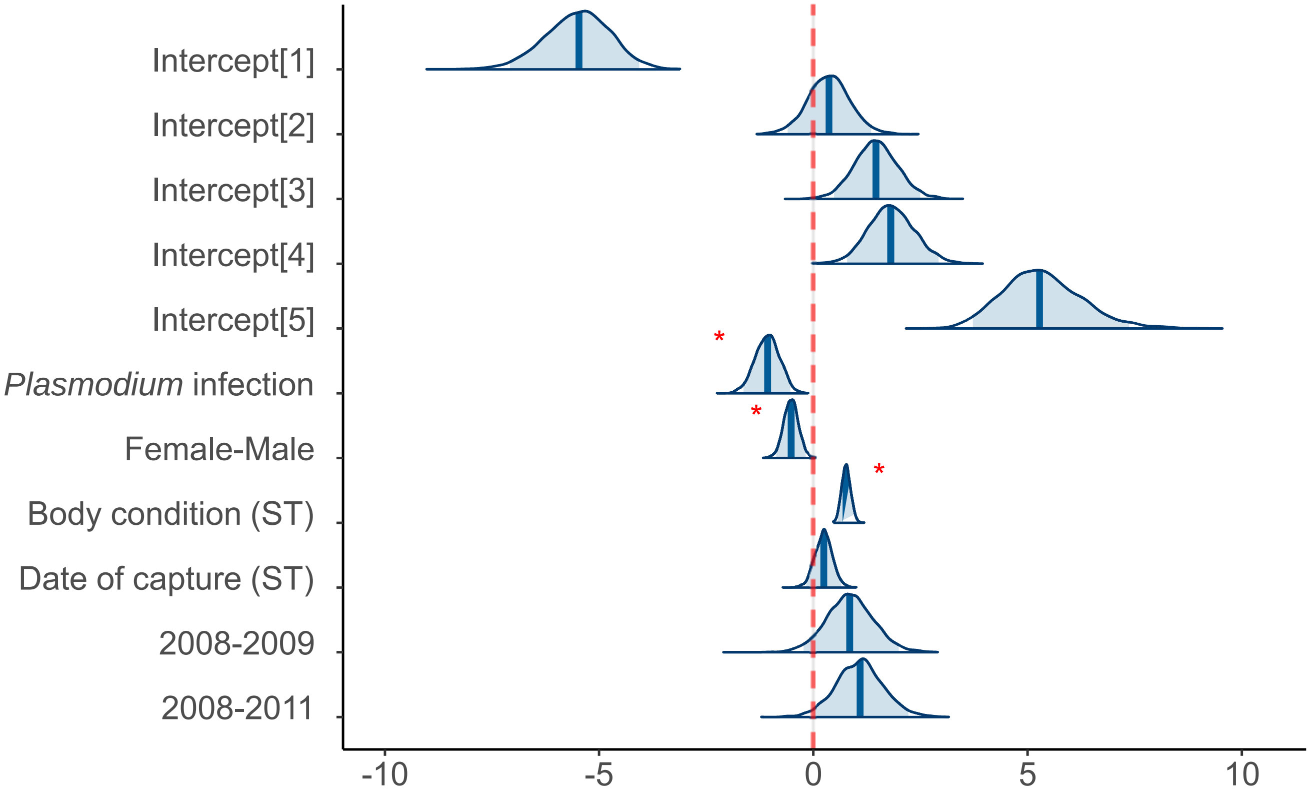

Electronic supplementary material A S3. Posterior distributions with medians and 95% confidence intervals for each parameter of the bayesian model assesing the influence of *Haemoproteus* status of infection on post-juvenile moult, controlling for year, sex, body condition and date of capture. We used * for significant comparisons with 95% BCI range values not overlapping zero and † for weak effects with 90%BCI range values not overlapping zero.

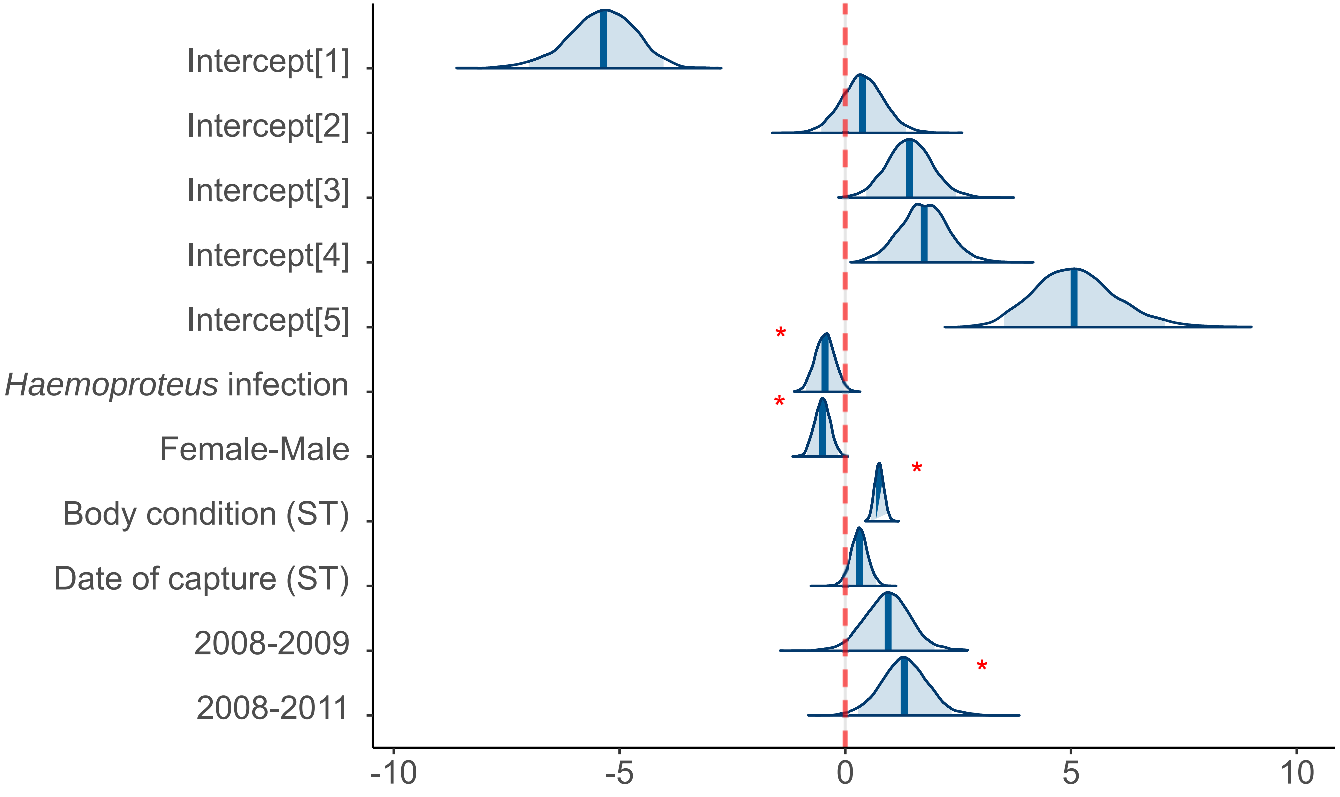

Electronic supplementary material A S4. Posterior distributions with medians and 95% confidence intervals for each parameter of the bayesian model assesing the influence of *Leucocytozoon* status of infection on post-juvenile moult, controlling for year, sex, body condition and date of capture. We used * for significant comparisons with 95% BCI range values not overlapping.

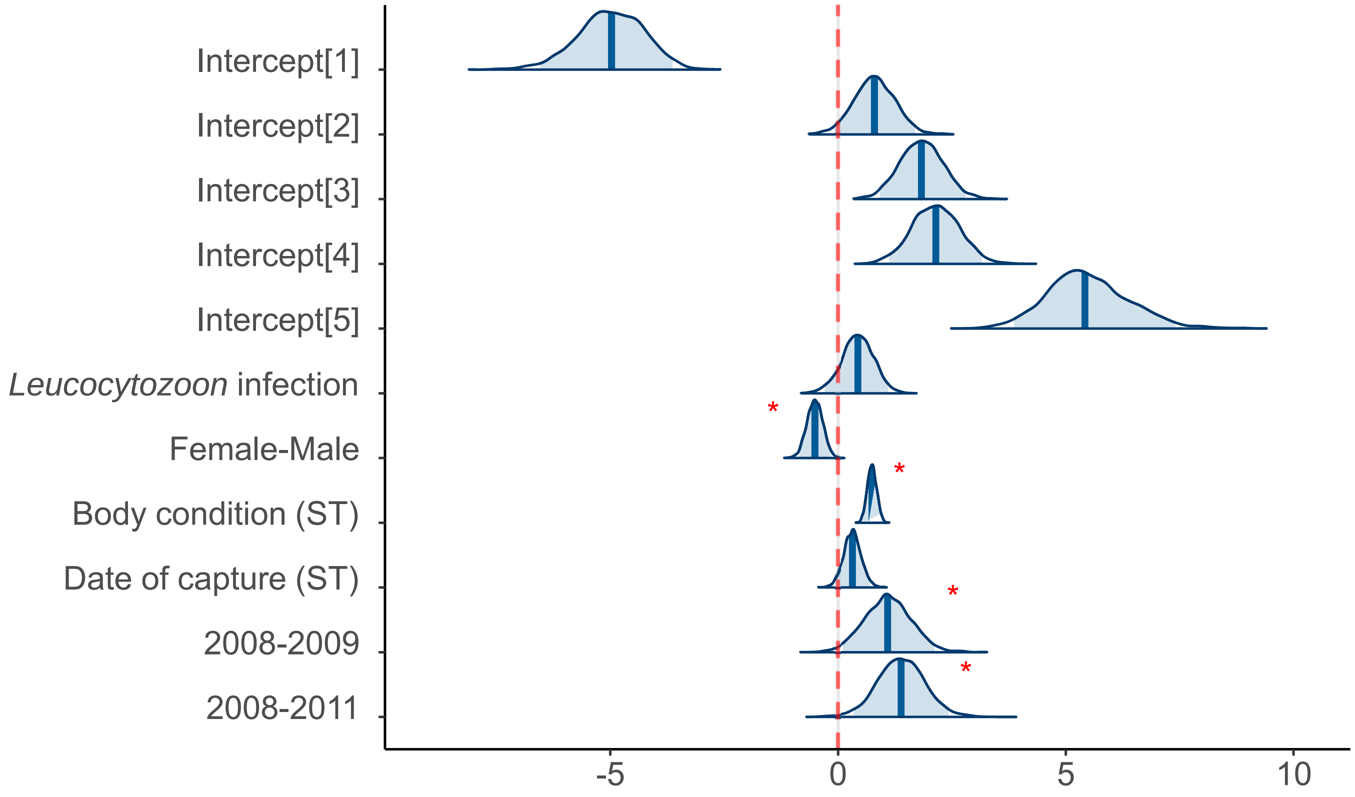

Electronic supplementary material A S5. Posterior distributions with medians and 95% confidence intervals for each parameter of the bayesian model assesing the influence of total parasite load on post-juvenile moult, controlling for year, sex, body condition and date of capture. We used * for significant comparisons with 95% BCI range values not overlapping zero and † for weak effects with 90%BCI range values not overlapping zero.

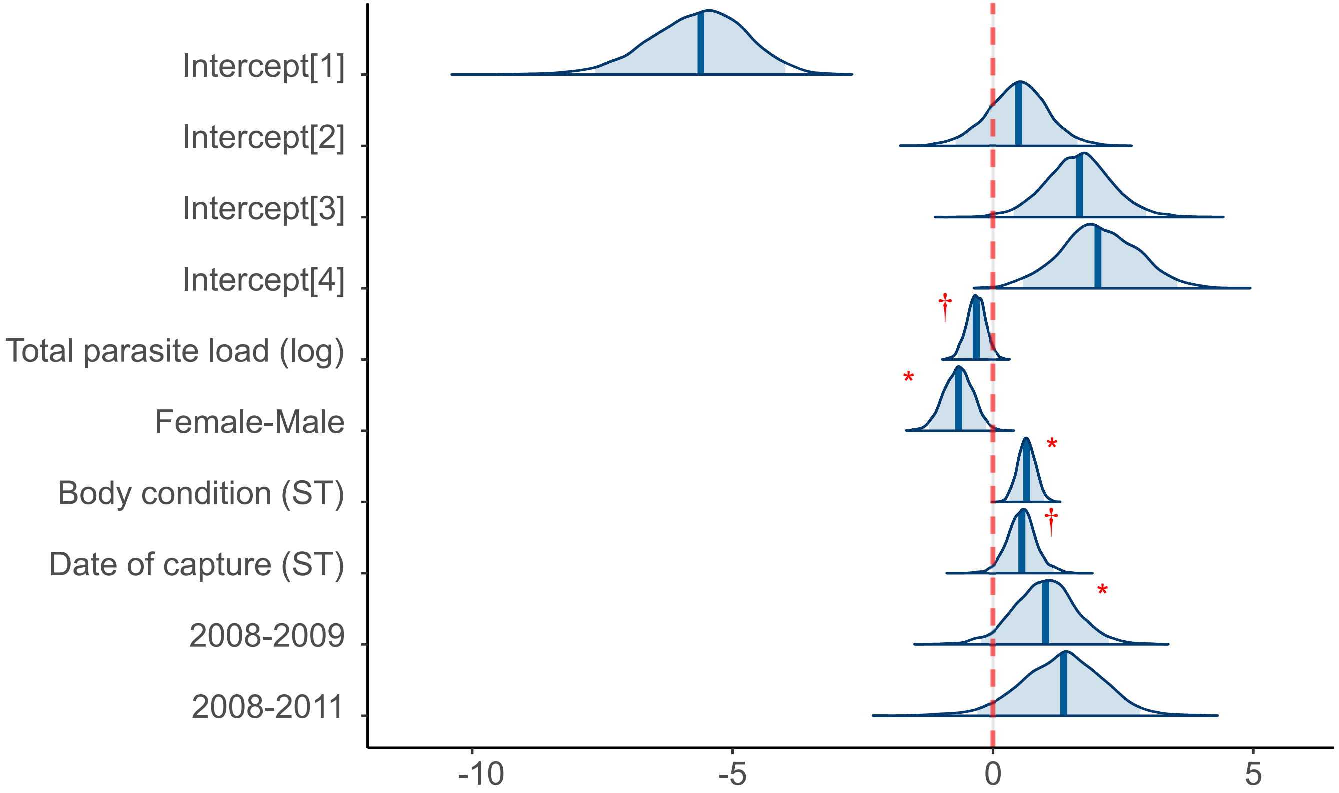

Electronic supplementary material A S6. Posterior distributions with medians and 95% confidence intervals for each parameter of the bayesian model assesing the influence of the presence of *Plasmodium* gametocytes on post-juvenile moult, controlling for sex, body condition and date of capture. We used * for significant comparisons with 95% BCI range values not overlapping zero and † for weak effects with 90%BCI range values not overlapping zero.

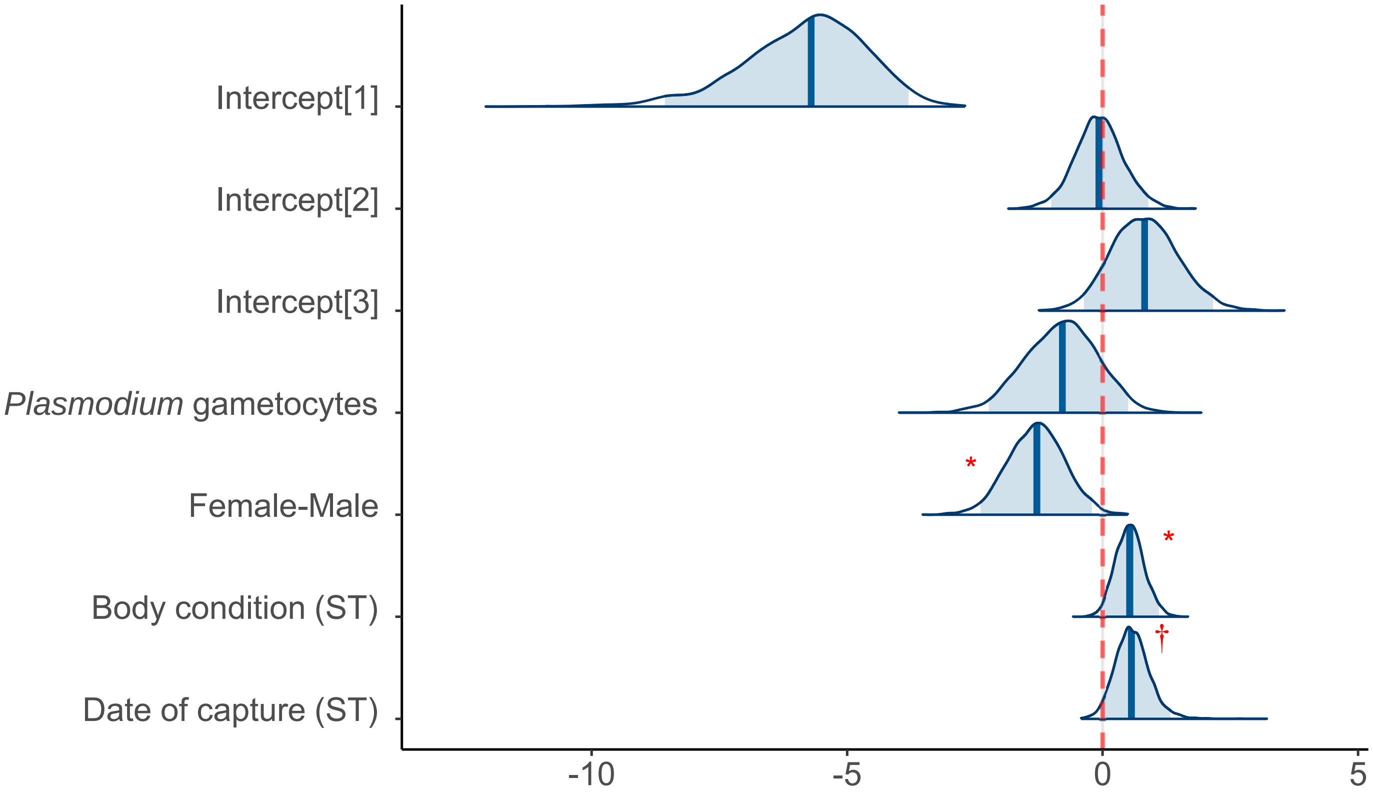

Electronic supplementary material A S7. Posterior distributions with medians and 95% confidence intervals for each parameter of the bayesian model assesing the influence of *Haemoproteus* parasite load on post-juvenile moult, controlling for year, sex, body condition and date of capture. We used * for significant comparisons with 95% BCI range values not overlapping zero.

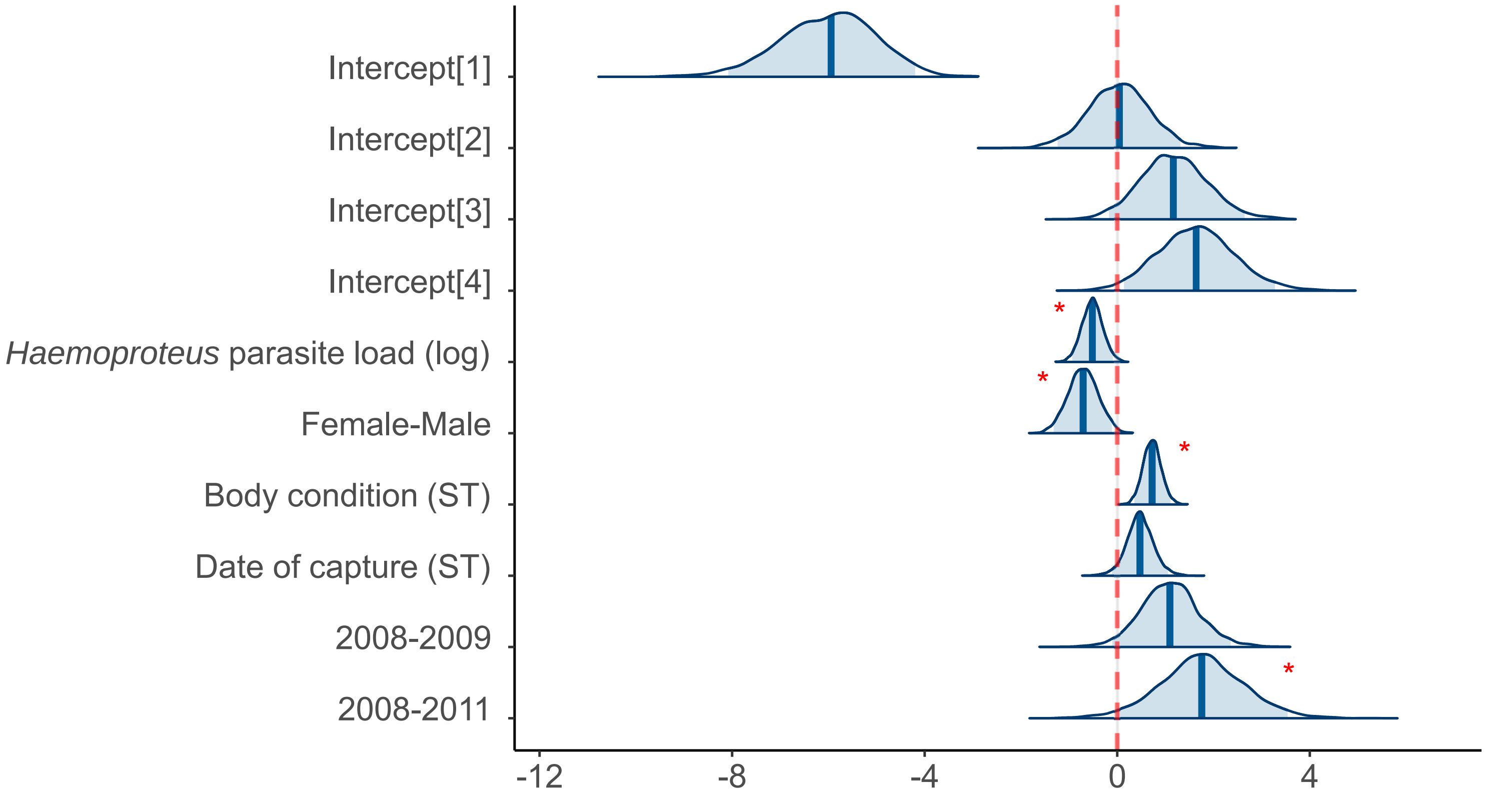

Electronic supplementary material A S8. Posterior distributions with medians and 95% confidence intervals for each parameter of the bayesian model assesing the influence of the presence of *Leucocytozoon* gametocytes on post-juvenile moult, controlling for sex, body condition and date of capture.

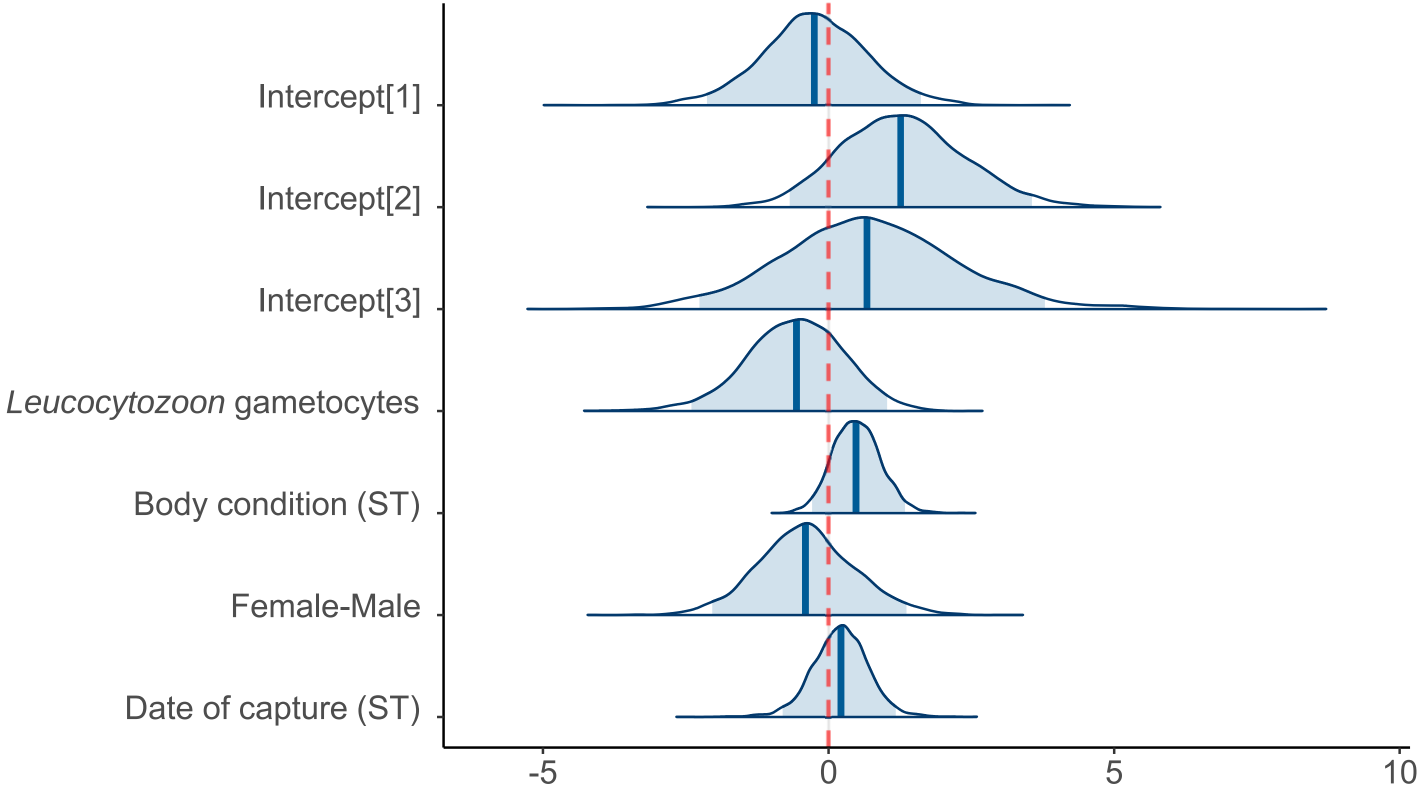
